# Insights into the linker domain in ABCB1/P-glycoprotein

**DOI:** 10.1101/2025.09.08.674578

**Authors:** Alessandro Barbieri, Nopnithi Thonghin, Richard F. Collins, Stephen M. Prince, Robert C. Ford, Hao Fan

**Affiliations:** School of Biological Sciences, Faculty of Biology Medicine and Health, The University of Manchester, Oxford Road, Manchester, M13 9PT, UK; Bioinformatics Institute (BII), Agency for Science, Technology, and Research (A*STAR), 30 Biopolis Street, #07-01 Matrix, Singapore 138671, Singapore; Department of Biology, Faculty of Science, Srinakharinwirot University, 114 Sukhumvit 23, Wattana District, Bangkok 10110, Thailand

**Keywords:** ABCB1, P-glycoprotein, Molecular Dynamics, Linker Region, Cryo-EM

## Abstract

ABCB1/P-glycoprotein(P-gp) is one of the best-known ABC transporters and its involvement in the clearance of chemotherapy agents in cancer cells can lead to multidrug resistance. The N- and C-terminal halves of P-gp are joined by a flexible linker region. Linker regions in ABC transporters often play pivotal roles (i.e. CFTR), however, there is no consensus on the importance of the role of the linker, the complete 3D-structure of which has not yet been determined. Prior studies on linker length showed major effects on P-gp function. In this work, we combined microsecond simulations (≈50µs) on mouse and human P-glycoprotein, cryo-EM and other computational methods to trace a profile of the linker region. Different experimentally verified conditions involving the linker region were simulated, analysed and compared. Attachment of the linker to the membrane was spontaneously and occasionally encountered via a recurrent series of amino-acids, despite random starting conformations, suggesting that the lipid bilayer might be involved in linker regulation. As carried out in previous experiments, shortening the linker could potentially disconnect the ATPase activity from the substrate transport by interfering with one ATP site while introducing a short partially-structured linker shows remarkable stability in accelerated MD. Finally, we investigated the linker region by positioning it in specific quadrants of the 3D space, with simulations suggesting 1) a possible influence on protein flexibility, 2) selective tightening and opening of the two halves of P-gp, and a 3) possible stabilization of the inward-facing/occluded-state. The interaction between the linker and ATPs was investigated using protein-protein docking, simulations and a re-refined cryo-EM map where medium-low resolution and weak Coulomb density was found between the nucleotide binding domains.

## Introduction

### ABCB1/P-glycoprotein and Linker region

Currently, 49 different ATP-binding cassette (ABC) proteins are known to exist in the human body and they are classified into seven subfamilies (from A to G). ABCB1, also known as P-glycoprotein (P-gp) or multidrug resistance protein 1 (MRD1) belongs to subfamily B of ABC transporters and it acts as a detoxifier of the body by extruding a large variety of chemotherapy agents from cells^1^. Overexpression or activation of P-gp can lead to a phenomenon called multidrug resistance (MDR) in cancer cells^2–5^. P-gp has been implicated in reduced bioavailability of therapeutic drugs in diseases like cancer and epilepsy^6,7^. In the human body, it is typically expressed in the apical membranes of the kidney, liver, pancreas, intestine^8^, placenta^9^ and capillary endothelial cells of the blood brain barrier^10,11^.

P-gp was first identified in 1971 in Chinese hamster ovary (CHO) cells selected for their resistance to colchicine^12^. Thereafter, roughly more than 28,000 studies have been carried out, including many structural biology studies. Although we have increased the number of structures representing distinct states of the transport cycle, none of these studies succeeded in the characterisation of the P-gp linker domain, and its exact functional role has never been completely clarified. Among more than three dozen human and mouse P-gp structures in the protein data bank (PDB), few of the ∼70 residues of the linker have been resolved, and these were mainly at both N-terminal and C-terminal extremities of the linker. The situation is similar in other ABC transporters, where parts of the linker are also missing from their structures. Examples of partly resolved linker regions in the eukaryotic ABCC family are ABCC1 (MRP1)^13^, ABCC2^14^, ABCC7(CFTR)^15^, and the yeast cadmium factor 1 protein (Ycf1p)^16^, in which several segments of the regulatory (R) region have been resolved by cryo-EM. In terms of functionality, the linker region in ABCC7/CFTR is known to be deeply involved in regulation. However, the CFTR has evolved into an ion channel from a transporter, making it an outlier among the other members of ABCC family. An example from the ABCB family comes from the inward-facing state of ABCB11 (PDB: 6LR0), which shows the presence of the linker inside its substrate binding cavity, similarly to ABCC2 (PDB: 8JX7) (**Supplementary Figure 1**). Whether this presence is part of the mechanism^17^ or simply a result of the affinity to peptides showed by many other ABC transporters is not known. Multiple functions or roles have been hypothesized for the ABC transporters linker regions^18^: ensuring their correct assembly, association with other proteins and function regulation. As a consequence, a question arises as to whether functions are mutually exchangeable or comparable with other ABC transporters, including P-glycoprotein.

Phosphorylation sites in human P-gp have been found at positions S661, S667, S671 and S683^23^ (**Figure 1**), with the last three strongly conserved in mammals. S667 and S671 are either targeted by protein kinase A (PKA) and protein kinase C (PKC), while S661 and S683 are a specific target for PKC or PKA, respectively. Corresponding studies on the effect of phosphorylation did not reach a consensus. Several mutagenesis studies of multiple serine residues found no impact on the ATPase and transport activity of ABCB1^24,25^, despite observations that suggested some correlations^18,26–30^. In contrast, reduced activity in WT and serine/threonine to aspartate mutants at lower concentrations of stimulatory drugs were found despite the maximum activity being unaltered^31^ and ATPase activity in ABCB1-expressing Sf9 insect cells was reduced by S671N mutation.

**Figure 1.**
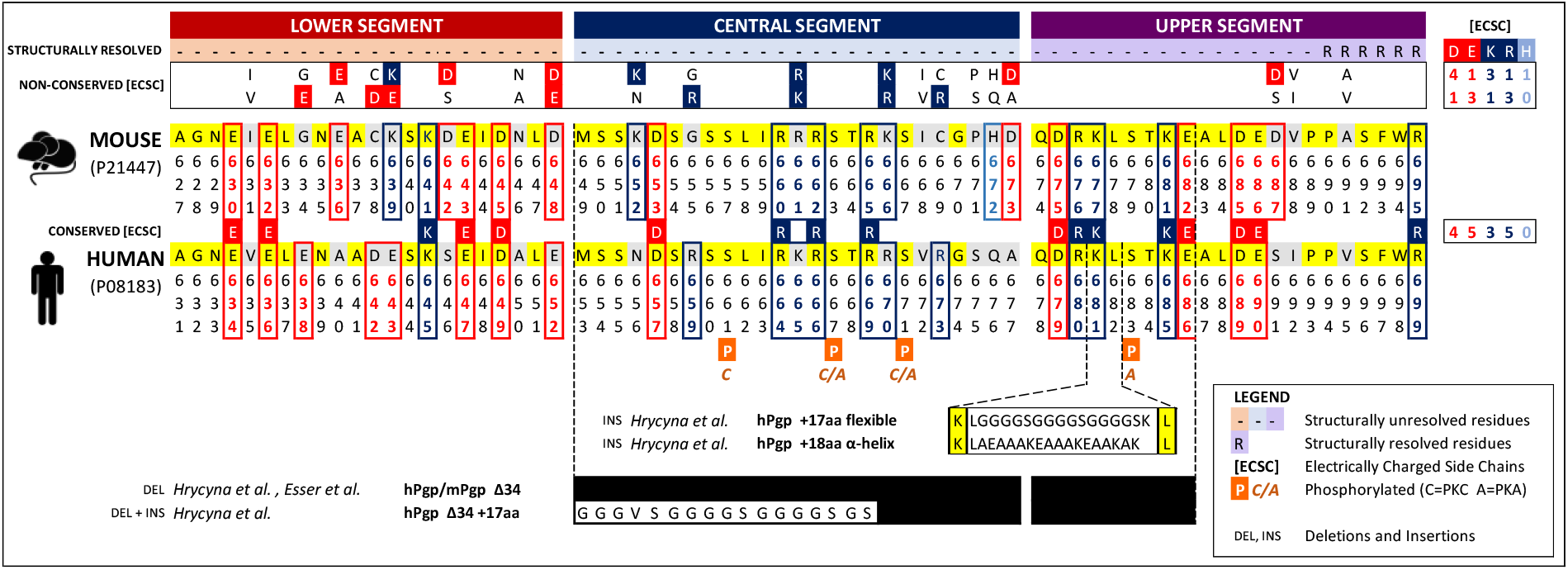
Sequence comparison between mouse and human linker regions. The linker sequence corresponds to ∼70 residues in mouse (UniProt entry P21447) and human (UniProt entry P08183) P-glycoprotein, with the sequence numeration shifted by four residues due to as many additional residues that are naturally present in the human extra-cellular loop 1 (ECL1). Electrically Charged Side Chains (ECSC) are coloured with blue and red boxes, while histidine residues were coloured in light blue. The linkers are subdivided into three distinct regions, coloured accordingly to the respective predominance of a specific electrically charged species. Conserved residues between species are reported in yellow boxes, non-conserved residues in grey boxes. Phosphorylation (orange boxes) sites are included for the human sequence with abbreviations (C=PKC and A=PKA). The conserved and non-conserved charged amino acids are counted on the right of this figure. On the bottom, references to prior studies were insertions/deletions of portions of the sequence were addressed. Black boxes represent the deleted parts.

Insertions and deletions of the linker region have been carried out in both mouse and human P-glycoprotein (**Figure 1**), in an attempt to correlate the linker length and composition to the regulation of ABCB1 function^32,33^. The deletion of 34 internal linker residues (653-686) in human P-glycoprotein^32^ (Δ34 hP-gp) did not suppress substrate nor ATP binding nor impeded the trafficking to the plasma membrane, but the basal and drug-stimulated ATPase activity was unexpectedly lost and so the ability of conferring drug resistance or transporting substrates. The removal of the corresponding 34 residues (649-682) in mouse P-glycoprotein (Δ34 mP-gp) affected drug transport activity *in vivo* or drug stimulation of ATPase activity *in vitro*, with both lost, but basal ATPase activity was still present.^34^ Interestingly, the Δ34 mP-gp structure was captured via X-ray crystallography but it neither permitted the identification of the ∼30 residues that remained, nor explained why shortening the linker affected P-glycoprotein function. The extension of the WT human linker via insertion of a flexible sequence of 17 residues did not alter the function or the conformation (detected by a conformation-specific antibody UIC2^32^); in contrast, the insertion of a hydrophobic, more defined helical structure composed by 18 residues provoked a loss of ABCB1 expression at the plasma membrane (**Figure 1**). However, after the combination of both deletion (34 residues) and insertion (17 neutral residues), the doubly-altered protein preserved 70-80% of the original activity.^34^

### Computational studies including the linker region of ABCB1

Numerous experimental studies on the linker region of ABCB1 have been carried out, as discussed above, however, almost all computational studies on P-glycoprotein are lacking the full-length linker. As exceptions, three distinct molecular dynamics (MD) studies on the 100-200 nanoseconds scale were reported in 2012, 2013 and 2020 by Ferreira, Bonito and collaborators on mouse/human P-glycoprotein^35–37^. Recently, human P-glycoprotein was studied with MD simulations on the 300-400 nanoseconds range and when the linker was modelled Elbahnsi^38^ and collaborators observed the cholesterol accessing the cavity from the membrane. Ground-breaking improvements in terms of hardware and GPU-enabled graphics cards^39^ were accomplished in the last decade, pushing the threshold of simulation length from nanoseconds to microseconds. In this work, we aimed to fill the knowledge gap on the missing linker region of P-glycoprotein by achieving microsecond (µs) MD simulations, cryo-EM refinement and docking calculations, ultimately comparing them with the existent experimental data.

## Results

### Linker Region: sequence analysis, predictions and modelling

The linker sequence in human and mouse P-glycoprotein corresponds to ∼70 residues, of which only ∼20 residues possess hydrophobic characteristics. Considering the sequence ranges of 627-695 in mouse and its corresponding 631-699 in human P-gp, the sequence identity of the two linkers is slightly lower than the global one between the species (71% vs 87%). This peptide conjoiner is highly charged (∼40%) in both human and mouse (**Figure 1**). The number of amino acids with electrically charged side chains (ECSC) at physiological pH (Arg, Asp, Glu, Lys) is almost identical in both species (25 human, 26 mouse). Interestingly, an almost neutral overall charge is present due to the internal equilibrium between positively and negatively charged residues (**Figure 1**). The small charge imbalance of basic residues present in the mouse linker could be potentially compensated by protonation of H672. The equilibrium is also preserved by separately considering the conserved (17) and non-conserved residues (8-10), where the sequence variations are not always consisting in a replacement of a charged residue with another. Interestingly, the number of non-conserved lysine and arginine residues is inverted between human and mouse P-gp (**Figure 1**) and almost exactly inverted in the case of glutamate and aspartate residues. Despite the overall charge balance, the charged residues are not equally spread, in fact, both linker sequences can be divided into three distinct regions (**Figure 1**): (1) a lower segment originating from NBD1 (631-652 human, 627-648 mouse) densely populated by negatively charged amino acids, (2) a central core segment (653-677 human, 649-673 mouse) populated by positively charged residues and, (3) a neutral upper segment (678-700 human, 674-696 mouse) reconnecting to the elbow helix at the start of TMD2.

Both sequences were analysed using PSIPRED, JNet and Robetta ab-initio to explore both secondary and tertiary structural features. These tools were used to detect regions that could possibly form secondary structure elements, similarly to what was observed in previous studies^35,36^. A specific segment within the central portion of the linker (according to the subdivision above) was predicted to form an α-helix by all methods (**Supplementary Figures 2**). However, in the case of Robetta *ab-initio*, the five models were generated by predicting only the central portion of the linker, since the two ends of the linker would have been unlikely to be predicted reflecting the natural distance that is present due to their attachment to NBD1 and TMD2. AlphaFold^40,41^ – as many PDB entries not immediately available at the dawn of our study – predicted no helical region in the centre of the human P-gp linker and two helical regions respectively in the first and third segments. In summary, some indications of helicity in the linker region emerged from these predictions, however due to the low confidence from predictions expected for these disordered regions, we adopted no structural features to be present in the linker, excepted when we directly tested a partially helical linker. Ultimately, for our MD simulations the missing parts in human or mouse P-glycoprotein were reconstituted through the loop modelling protocol within UCSF Modeller. For each system, corresponding to a specific experimentally verified condition, 2500 different linkers were generated with the top ranked models chosen for simulations (see Methods).

### MD simulations on P-glycoprotein with linker reconstructed

Our investigations on the linker region and its possible effects on ABCB1/P-glycoprotein were conducted by initially considering a series of conditions that were already experimentally tested through prior mutagenesis and functional studies. The following systems were therefore considered: 1) human and mouse WT, 2) extended and shortened linker (human, mouse) and finally a 3) partially structured linker (**Table 1**). In a second set of simulations, we also probed different conformational states of the P-glycoprotein, corresponding to different phases of the cycle. Each simulation condition was conducted in multiple replicates of minimum length of 1 microsecond (μs) to give indications of reproducibility.

**Table 2.** Full panel of the MD simulations (simulation time expressed in microseconds, µs). Legend: *Gaussian accelerated MD, **Revised cryo-EM density, (HM) Homology Modelling.

| System | Organism | STATE | PDB | RE 1 | RE 2 | RE 3 | RE 4 |
| --- | --- | --- | --- | --- | --- | --- | --- |
| WT IF | Human | IF closed | 6qex | 1.0 | 1.0 | 1.0 | 3.0 |
| OF | Human | OF | 6c0v | 1.0 | 1.0 | 1.0 | 3.0 |
| SS (α-helix) | Human | IF closed | 6qex | 1.0 | 1.0 | 1.0* | 1.0* |
| Length (+17aa flexible) | Human | IF closed | 6qex | 1.0 | 1.0 |  |  |
| Length (+18 aa helical) | Human | IF closed | 6qex | AF |  |  |  |
| Length (Δ34) | Mouse | IF wide-open | 4xwk | 1.0 | 1.0 | | |
| Length (Δ34) | Mouse | IF Δ34 | 5koy | | | 1.0 | 1.0 |
| WT IF | Mouse | IF wide-open | 4xwk | 1.0 | 1.0 | 1.0 |  |
| Conformation (pos.1) | Human | IF closed | 6qex | 0.25 | 0.25 | 0.25 |  |
| Conformation (pos.2) | Human | IF closed | 6qex | 0.25 | 0.25 |  |  |
| " | Human | IF wide-open | 4xwk (HM) |  |  | 0.25 | 0.25 |
| Conformation (pos.3) | Human | IF closed | 6qex | 0.25 |  |  |  |
| " | Human | IF wide-open | 4xwk (HM) |  | 0.25 | 0.25 |  |
| Conformation (pos.4) | Human | IF closed | 6qex | 0.25 | 0.25 |  |  |
| " | Human | IF wide-open | 4xwk (HM) |  |  | 0.25 | 0.25 |
| " | Mouse | IF wide-open | 6q81** | 0.25 | 0.25 | 0.25 | 0.25 |
| Conformation (pos.5) | Human | IF wide-open | 4xwk (HM) | 0.25 | 0.25 | 0.25 | 0.25 |
| Conformation (pos.6) | Human | IF wide-open | 4xwk (HM) | 0.25 | 0.25 | 0.25 |  |
| Conformation (pos.7) | Human | IF wide-open | 4xwk (HM) | 0.25 | 0.25 |  |  |

The possible helicity of the P-gp linker region is implied not only by computational predictions but also by HDX-MS data^44^, therefore we tested a partially structured linker region in WT human P-gp. Using Modeller, we generated several human P-gp models with a α-helix modelled in the central linker segment, where most of the predictions converge. Four simulations with different linker starting conformations were carried out, including two replicas where we employed an enhanced sampling protocol with the usage of Gaussian accelerated MD (GaMD)^45^. In the unbiased replicas, the structured linker showed no sign of disruption in its α-helix, neither in the single case where a spontaneous attachment of the linker to the lipid bilayer occurred (**Supplementary Figure 5a**). Instead, we observed a stabilization of the conformation and an increased helicity in other portions of the linker. To assess the stability of the linker and to enhance the conformational sampling, we employed Gaussian accelerated MD (GaMD)^45^ within the Amber package, following a previous protocol^46^. Remarkably, after 1 µs of enhanced sampling, only few residues at the ends of the α-helix lost their helicity and the initial structural organization. Interestingly, the presence of the helix in the simulations reduced the number of residues involved in forming disordered tangles within the linker region (**Supplementary Figure 3**).

### MD simulations of P-glycoprotein with shortened/extended linker

As evidenced from the P-glycoprotein background, the length and composition of the linker were experimentally modified through insertions, deletions or both^47^. We computationally investigated the variation in the linker length, starting with the extension. The linker sequence was modelled based on the WT IF closed conformation (PDB 6qex) and using the same Modeller protocol as described in the Methods. Two simulations (1 µs each) of human P-gp with a 17-residue flexible peptide inserted in between residues 681-682 were carried out. From visual inspection and the analyses of trajectories, no significant changes on the conformation of NBDs and TMDs were observed (**Supplementary Figure 6c**). Interestingly, the RMSD of the linker, normally reaching a plateau after 100 ns in human WT simulations, was shifted to around 300-400 ns. Nevertheless, the TMDs and NBDs stabilized at a final RMSD of 2-3 Å, as seen for the WT counterpart (**Supplementary Figure**). In terms of the appearance of secondary structure in the linker, we observed some helical formation in the first segment, whilst no structural element was observed in the other parts, except for a transient 3_10_-helix (**Supplementary Figure 3**). Due to its flexibility, the extended linker appeared to agglomerate into a single mass.

The insertion of an 18-amino-acid peptide sequence expected to form a helix in human P-gp resulted in an unstable protein, not reaching the plasma membrane^47^. Consequently, we opted to not carry out MD simulations of the system as it would not be consistent with experiments. However, with AlphaFold^41^, the extended linker was predicted to form a much longer helix (**Supplementary Figure 6a**,**b**) than expected. The additional helix-forming residues in the wild-type linker were already predicted to be more likely to adopt a helical structure (**Supplementary Figure 2**) although at low level of confidence.

Lastly, we focused on the experiments where the linker was shortened by 34 residues, removing the entire central segment and a part of the third segment (**Figure 1**). Unlike for the human homolog, an X-ray crystal structure (PDB: 5koy) was obtained for the mouse P-gp (Δ34 mP-gp). Interestingly, this structure showed a substantial reduction (Å) in the level of opening of the NBDs compared to the wild-type protein (Å)(**Supplementary Figure 7a**). Additionally, a single molecule of ATP was found to be bound to NBD1 in this structure (albeit the absence of Mg ions in the crystallization conditions). According to Esser and collaborators^34^, the linker-shortened mutant protein had a slightly elevated basal ATPase activity but it had lost the drug-stimulated ATPase activity *in vitro* and the ability to transport substrates *in vivo*. Microsecond simulations for Δ34 mP-gp^5KOY^ were carried out by modelling the original crystal structures (PDB 5koy) with the starting conformations of the linker either located between the two NBDs or on the outside of them (with limitations due to its reduced length). As a comparison, simulations of the wild-type mouse P-glycoprotein were included as negative control. In contrast to results for the extended linker, the RMSD plateau of the linker occurred after 100 ns, similarly to the human and mouse WT simulations.

Perhaps surprisingly, Δ34 mP-gp^5KOY^ simulations showed a further decrease of the already reduced NBDs distance and a substantial reduction of the substrate binding cavity across the whole production. Significantly, no secondary structural elements were detected during the simulations, compared to the situation in the WT and the extended linker simulations. However, the shortened linker did occasionally display some tendency to collapse in on itself, similarly to observations for WT and extended linker simulations.

If the shortened linker was modelled on the mouse IF wide open state (PDBs 4xwk/5kpj), after several hundreds of nanoseconds, the two Δ34 mP-gp^4XWK^ replicas converged to the experimental X-ray structure for the shortened linker (PDB 5koy) in terms of the opening level of the two halves (**Supplementary Figure 7b**,**c**,**d**,**g**). This reduction of the distance between the NBDs was not observed in the WT mouse P-gp control simulations (**Supplementary Figure**). Furthermore, in both Δ34 mP-gp^4XWK^ simulations, we observed the upward (towards the membrane) tilt of the NBD1-terminal helix as sort of compensation mechanism to the reduction in length (**Supplementary Figure 7c**,**d**). The reduced length impeded structural re-arrangement as instead observed in the wild-type simulations (**Supplementary Figures 2 and 3**). In Δ34 mP-gp^4XWK^ and Δ34 mP-gp^5KOY^, the shortened linker occupied a position between the two NBDs (**Supplementary Figure 7g**), in particular in an area adjacent to the nucleotide-binding site 2 (NBS2).

### Linker conformations in the 3D space

During this investigation, we focused on exploring different linker conformations to study several experimentally verified conditions. A key thinking that emerged from simulations is whether the possible effect exerted by the linker to the whole protein are depending on the linker localisation upon reaching the RMSD plateau. The importance of the localisation has arisen from distinct simulations showing the linker remaining spontaneously and steadily attached to the lipid bilayer or closer to specific regions of the protein. Throughout a selective modelling process combining – when necessary – UCSF Modeller with Isolde and minimization (see Methods), we positioned the linker in specific quadrants of the cytoplasmic 3D space not occupied by P-glycoprotein and potentially reachable by the linker region (**Figure 4**). The subdivision into eight quadrants was made after considering that the linker could potentially 1) lie in the water medium outside or between the two halves of P-glycoprotein, 2) occupy different heights or 3) flank the protein at each side. Since the two ends of the linker are attached to NBD1 (quadrant 4) and TMD2(quadrant 1), the threshold of how many residues needs to be lying into a specific quadrant to assume the linker is in a focused position was established.

**Figure 4.**
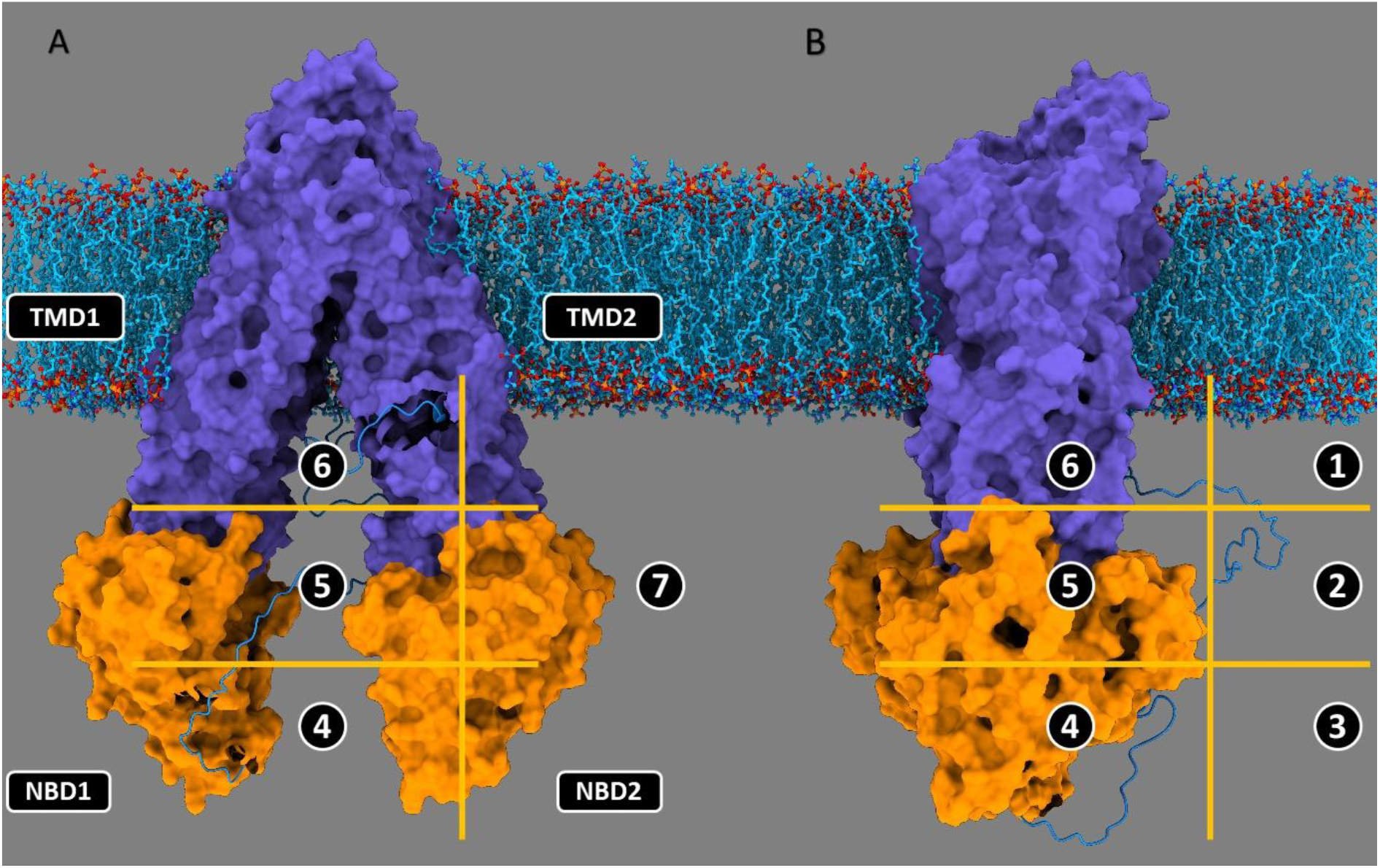
Key quadrants (1-7) for testing possible effects exerted by the linker depending on its localisation. Upon targeted modelling capable of locating the linker predominantly in specific quadrants, short simulations were conducted to verify the linker mobility and effects upon the interaction with different portion of the protein or with the plasma membrane. Positions in quadrant 1 to 3 are external, while 4 to 6 lie in between the two halves of P-gp. Positions in quadrant 7 and 8 flanks NBD2/TMD2 and NBD1/TMD1, respectively. **A)** Back view of human P-gp with reconstructed linker derived from homology modelling of mouse IF wide-open state (PDB 4XWK as template). **B)** Lateral view of human P-gp with reconstructed linker.

Quadrants 1 to 3 correspond to the linker lying outside of the two halves of P-gp (**Figure 4**), regardless of the level of opening of NBDs, thus both IF wide-open and IF closed conformational states were employed (PDBs 6qex and 4xwk). Since the close distance between NBDs in the human IF closed (PDB 6qex) would have not been compatible for studying a focused linker in quadrants 4-6, we generated the linker using the mouse IF wide-open conformation (PDB 4XWK) as template for homology modelling. As of today, a human IF wide-open state (PDB 9CTF) is present in the protein data bank and this IF wide-open conformation is perfectly matching the mouse P-gp employed in our studies (**Supplementary Figure**).

### Cryo-EM of mouse P-glycoprotein and linker region

Based on current entries in the protein data bank, no deposited structure shows the missing linker, except for several additional residues^51,52^. Elongated additional densities were located at the expected nucleotide-binding site 1 (NBS1) in the cryo-EM map EMD-4391 (PDB 6Q81) from our previous work^52^, although the low resolution 7.9Å did not allow the modelling.

## Discussion

Possible roles of linker regions in ABC transporters were studied due to their potentially high structure-to-function correlation.

While this occurred especially for members of the ABCC subfamily, the linker region of P-glycoprotein was also extensively studied during the past decades, despite a consensus on its role was not reached. The aim of this work was to provide extensive computational support to a broad range of previously reported experiments and to fill the existent knowledge gap. This included investigating various conditions as well as the impact of shortening/increasing the length of the linker. Currently, there is no structure available of the linker region for mouse and human P-gp, which share an 87% sequence identity. The only possible traces of the linker were observed in our previous electron-density map^59^ of mouse P-glycoprotein, whose resolution was substantially improved in this work. We could subdivide the linker into (1) lower, (2) central and (3) upper segments (**Figure 1**). The upper segment contains an equal number of residues with electrically charged side chains (ECSC) and it reconnects the linker to the elbow helix at the start of TMD2. The situation drastically changes for the other two segments: a lower segment, in direct prosecution of the final helix of NBD1, mainly composed by acidic residues and, in contrast, the dominance of basic residues showed in the central segment. Interestingly, the charge balance is preserved by considering each individual species or subsets such as conserved or non-conserved residues between the two.

The charge preservation and the specific distribution of basic and acidic residues in the lower and central segments is also present in mouse **ABCB4** (or MDR3 encoded by *mdr2*), a phosphatidylcholine flippase, whose linker was proven to be exchangeable with the corresponding of mouse P-gp with no functional variation of activity (**Supplementary Figure 1**)^60^.

The missing linkers in human and mouse P-glycoprotein were reconstituted using the loop modelling protocol within UCSF Modeller (see Methods), adopting no secondary structure elements and sampling different linear linker conformations. As templates, we employed two distinct inward-facing (IF) conformations: human IF closed state (PDB 6QEX) ^61^ and mouse IF wide-open state (PDB 4XWK), which was lately found to be perfectly matching the first IF wide-open ever deposited^62^. This alternance allowed to study the linker outside or in between the two halves of P-glycoprotein. We carried out a series of microsecond simulations on both human and mouse P-glycoprotein to mimic different experimentally tested conditions. A commonality encountered in all simulations is the stabilization of the linker in terms of RMSD value after an average of 100 ns (**Supplementary Figure 12)**, possibly to the rapid achievement of a conformational equilibrium or due to local minima, whose ‘trap’ could be potentially overcome with longer simulations.

To test the various predictions from PSIPRED, JNet and Robetta *ab initio*, we generated a series of partially structured linkers with an α-helix present in the central core. Gaussian accelerated MD (GaMD) simulations were employed to test if the integrity of the helix could be perturbated during enhanced sampling. Both GaMD simulations showed a remarkable stability of the α-helix after one microsecond each (**Supplementary Figures 2 and**).

Further helical arrangements in other linker segments were observed in all simulations, in accordance with computational predictions and with HDX-MD data^66^. (**Supplementary Figures 2 and 3**). During the simulations, the linker region generally underwent local collapse and coiling, adopting a compact globular-like conformation. This phenomenon seems to be more associated with the free and untangled starting conformations generally employed, while adding a 10-residues long α-helix to the linker central segment apparently reduced the formation of secondary structure elements (**Supplementary Figure 2**), at least in the microsecond timescale.

Hrycyna and collaborators performed a series of experiments to show the effects of extending, shortening or changing the composition of the linker in human and mouse P-gp (**Figure 1**).^32,67^ Since these systems were never reproduced using computational tools, several modelling solutions and simulations were developed. Notably, the extended linker with a peptide supposed to adopt a helical structure was predicted by AlphaFold3^41^to form a much longer helix than expected, combining the 18 residues from the peptide with several wild-type residues of the third segment, already predicted with low-confidence to be prone to helicity in the wild-type protein (**Supplementary Figures 2 and 6a**,**b**). If verified, this hypothetical longer helix could potentially interfere with the proper folding, as originally hypothesized by Hrycyna and collaborators^68^ and explain the absence of protein trafficking to the plasma membrane.

## Methods

### Homology modelling and secondary structure prediction

Homology Modelling of the linker was carried out by employing UCSF Modeller 9.23^78^ and where necessary, the protocol was implemented to contain two molecules of ATPs bound to NBDs. An amount of 2500 different linkers per model were generated with DOPE and MOLPDF employed to score the models. Final models were chosen on the basis of a general consensus of top ranked results for both scores, following a common approach for every system. Secondary structure predictions were carried out via JNet, PSIPRED while to assess if the central core of P-gp linker (mouse 664-673, human 668-677) could form any ordered structure Ab-initio ROBETTA was employed. Linkers were thus generated without secondary structure restraints in order not to bias the simulations. To explore those regions not covered by the conventional homology modelling, we employed the Isolde plugin within UCSF ChimeraX. The Interactive molecular dynamics panel of Isolde was employed to dynamically re-orient the linkers by iteratively guiding them towards targeted areas of the 3D space allowing to generate novel random conformations.

### System preparation and molecular dynamics

MD simulations were carried out using the AMBER package, version 20 with AmberTools21. Systems preparation was achieved by using the CHARMMGUI web interface for membrane embedding, solvation and balancing of salt concentration. Classic MD simulations of P-glycoprotein were carried out following a previous setup for membrane proteins^79^ with necessary adjustments. All systems were placed in a rectangular water box with periodic boundary conditions, employing TIP3P explicit solvent model and a concentration of 0.15M of KCl. An appropriate number of necessary counterions were added to neutralize the overall charge of the system. Amber FF14SB was employed as force field with the particle mesh Ewald method for long-range electrostatic interactions and a cut-off of 10 Å for the nonbonded interactions. Time step was set at 2.0 femtosecond and the SHAKE algorithm was applied. Each system was energy minimized using steepest descent algorithm followed by conjugate gradient and the temperature was gradually increased in two steps from 0 to 300K. Temperature and pressure were maintained constant using anisotropic pressure scaling and Langevin thermostat. Equilibration was staged in various steps, with a gradual decrease of restraints until final relaxation. To process and analyse all trajectories, various tools were employed: CPTRAJ, VMD and UCSF Chimera.

## Supporting information

SuppInfo

## Acknowledgments

All material contained in this paper derives from the Ph.D. thesis of Dr Alessandro Barbieri (The University of Manchester, A*STAR Singapore). We would like to thank both Hao Fan’s and Robert C. Ford’s research groups for their support and interesting discussions. The authors would like to acknowledge the assistance given by Research IT and the use of the Computational Shared Facility at The University of Manchester. A special thank for their kindness, speed and availability. These computational studies were also conducted using the computational facilities of the Bioinformatics Institute of A*STAR in Singapore and of the Singapore’s National SuperComputing Center (NSCC). A special thank for their kindness, speed and availability. We would like to thank the people of the Amber-Hub website for keeping updated the scripting support for CPPTRAJ.

