## Supplementary material for "Insights into the linker domain in ABCB1/P-glycoprotein": SuppInfo

### Table of Contents

**Figure S1.** Linker regions in ABCB1, ABCB4, ABCB11 and ABCC2.

**Figure S2.** Secondary structure predictions of human P-glycoprotein and DSSP analysis of several simulations testing experimentally verified conditions.

**Figure S3.** DSSP analysis of human and mouse P-glycoprotein simulations testing different conditions.

**Figure S6.** Human P-glycoprotein with extended linkers according to the experimental work.

**Figure S7.** Models and simulations of P-glycoprotein with shortened linker (delta653-686).

**Figure S12**. Root mean square deviation (RMSD) of the linker regions in the human P-glycoprotein simulations.

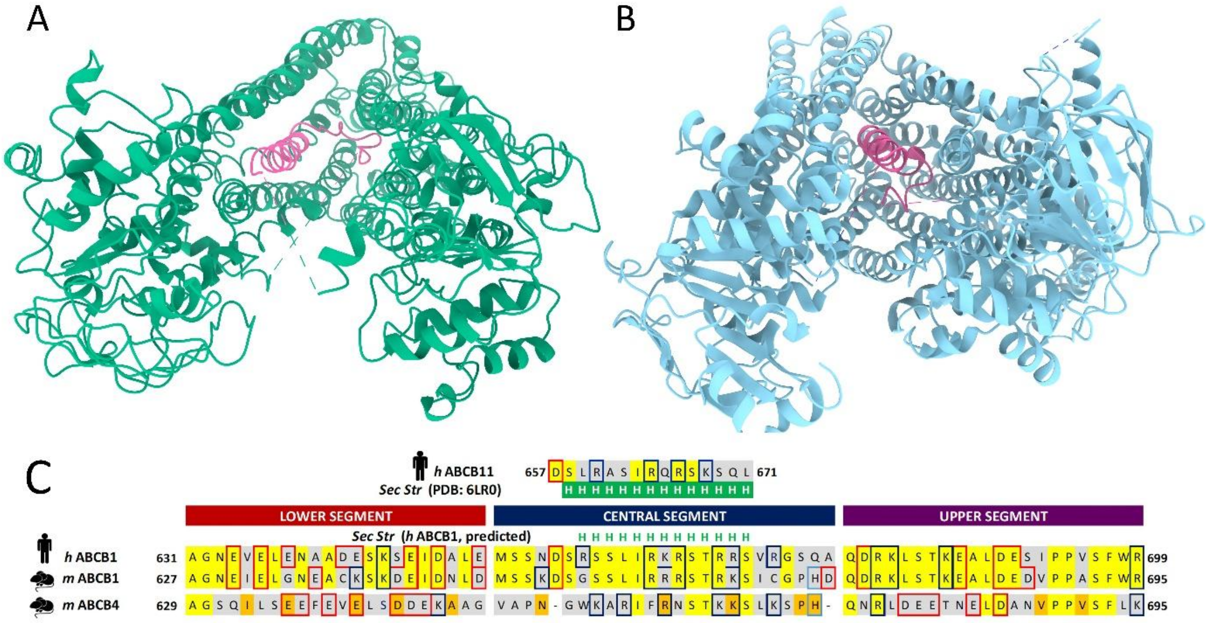


**Figure S1.** Linker regions in ABCB1, ABCB4, ABCB11 and ABCC2. **A)** ABCB11 (PDB: 6LR0) is colored in green and the α-helix formed inside the cavity is 14 residues long. **B)** ABCC2 (PDB: 8JX7) is colored in light blue and the α-helix formed inside the cavity is 16 residues long. Both ABCB11 and ABCC2 were determined in inward-facing (IF) state by single-particle analysis in cryo-EM. **C)** Sequence alignment and comparison of linker regions in mouse/human ABCB1 and mouse ABCB4. These sequences were subdivided into three regions following our classification (see Results). Conserved residues are represented in yellow boxes while grey boxes are reserved to the non-conserved ones. Amino acids with electrically charged side chains (ECSC) are colored accordingly. The secondary structure of ABCB11 is reported for comparison with predictions of human ABCB1.

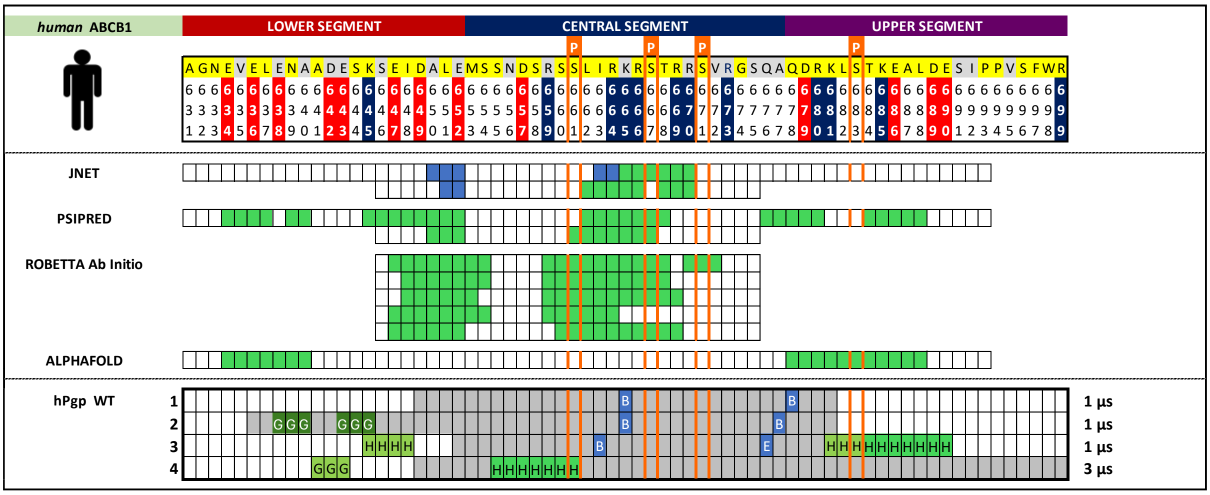


**Figure S2.** Secondary structure predictions of human P-glycoprotein and DSSP analysis of several simulations testing experimentally verified conditions.

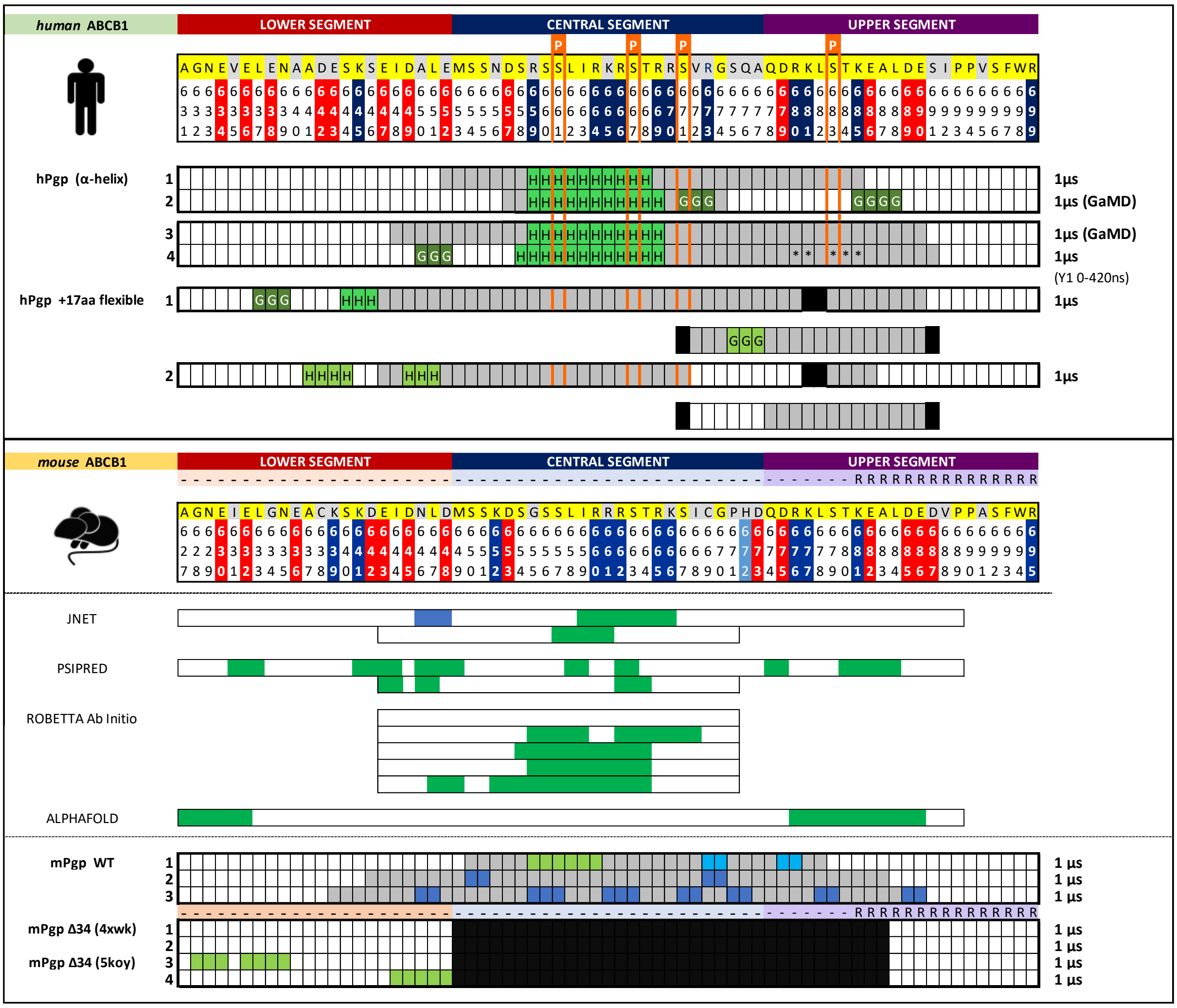


**Figure S3.** DSSP analysis of human and mouse P-glycoprotein simulations testing different conditions: extension and deletion of the linker, the addition of an α-helix within the linker and mouse WT P-gp control simulations were carried out.

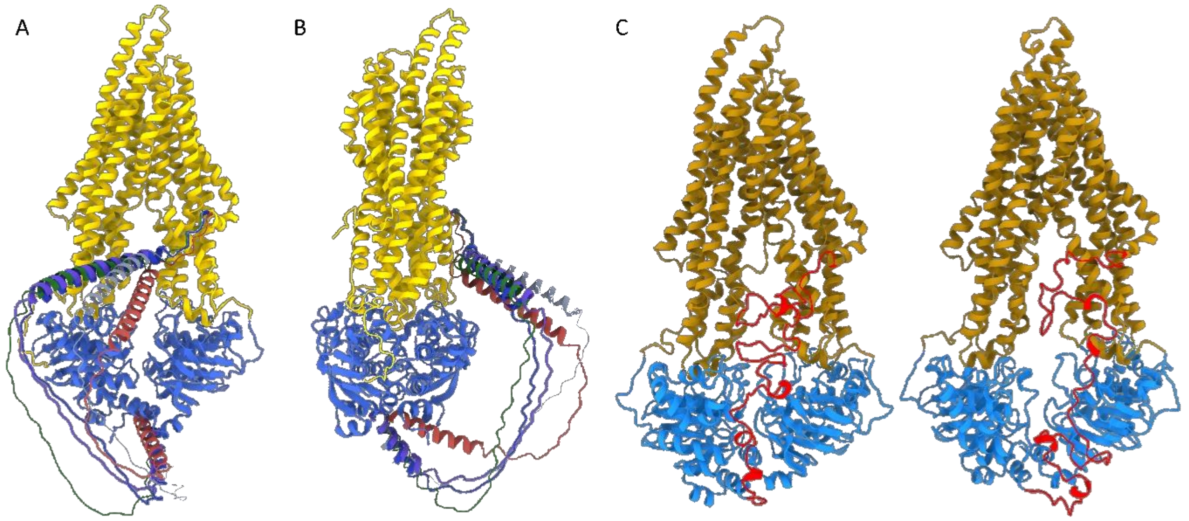


**Figure S6.** Human P-glycoprotein with extended linkers according to the experimental work. **A,B)** Back view and lateral view of AlphaFold3 models of human P-glycoprotein with an 18 aa peptide insertion with a predicted alpha-helical structure. **C)** MD simulations of human P-glycoprotein with 17 aa flexible peptide sequence inserted in between positions 681-682. Representations taken after clustering with k-means of the two replicas of the extended linker with a flexible sequence. These models or simulations are based upon the experiments conducted by Hrycyna and collaborators[1] where two different sequences were added in between amino acids K681 and L682. The hypothetical prediction of AlphaFold3 seems to combine the 18 a.a. helical peptide with some native residues – predicted as helix by AlphaFold3 in human WT P-gp – in a unique secondary structure element.

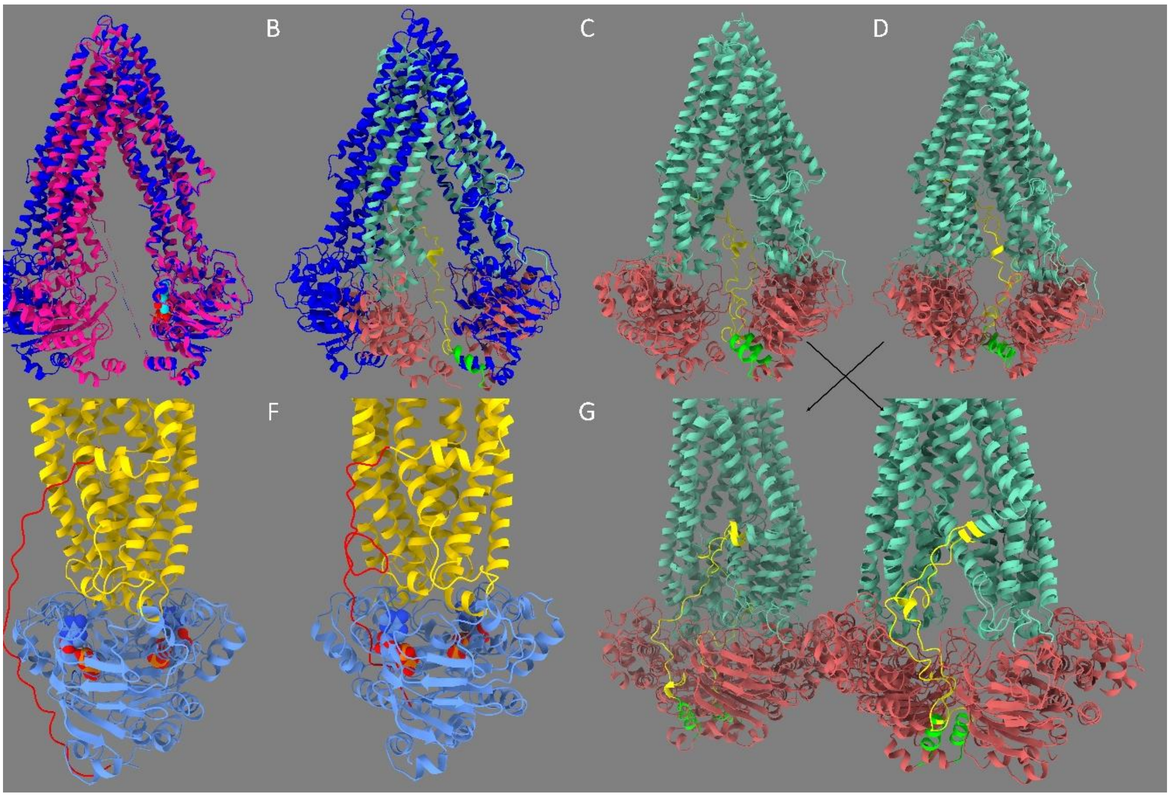


**Figure S7.** Models and simulations of P-glycoprotein with shortened linker (delta653-686). **A)** A comparison between mouse WT P-gp (PDB 4xwk, blue) and its relative simulation with shortened linker (yellow). TMDs (aquamarine blue) and NBDs (red) in the simulations naturally come closer to each other after hundreds of nanoseconds. **B)** Differences in terms of opening level of the NBDs between PDBs 4xwk (blue) and 5koy (delta653-686, deep pink) are showed. PDB 5koy presents an ATP molecule on NBD1 (cyan, sphere format). **C,D)** Simulations respectively starting from PDB 4xwk and from PDB 5koy, converging to a final level of opening of the NBDs similar to the X-ray structure of Δ34-mouse P-gp. The differences of length with WT P-gp seemed to have been compensated by the terminal helix of NBD1 (lime green) preceding the linker, which adjusts itself and points upward to reduce the gap, naturally remedying to the shortened length. Observed a disengagement of the NBD2 in one replica, event also by other groups. **E,F)** Representation of hypothetical outward-facing P-gp models with shortened linker. TMD domains are colored in yellow, NBD domains in cornflower blue. ATP molecules are represented in sphere format, the linker is colored in red. **E)** In this homology model, the shortened linker is just sufficient to not remain trapped in between the two halves during the ATP-dimerization that leads to the outward-facing state (OF). **F)** A hypothetical representation of the linker remaining trapped, hypothetically interfering with the correct ATPase activity and ATP dimerization. **G)** Back view of mouse P-gp simulations with shortened linker (delta653-686) with a focus of the linker occupancy. In all replicas, the shortened linker lies in the middle of the two wings, it interacts with the protein and potentially could interfere with the occupation of one nucleotide-binding site (NBS2). ATP is not showed in this representation. On the left of panel **G)** simulations starting from PDB 5koy, while on the right, simulations starting from PDB 4xwk.

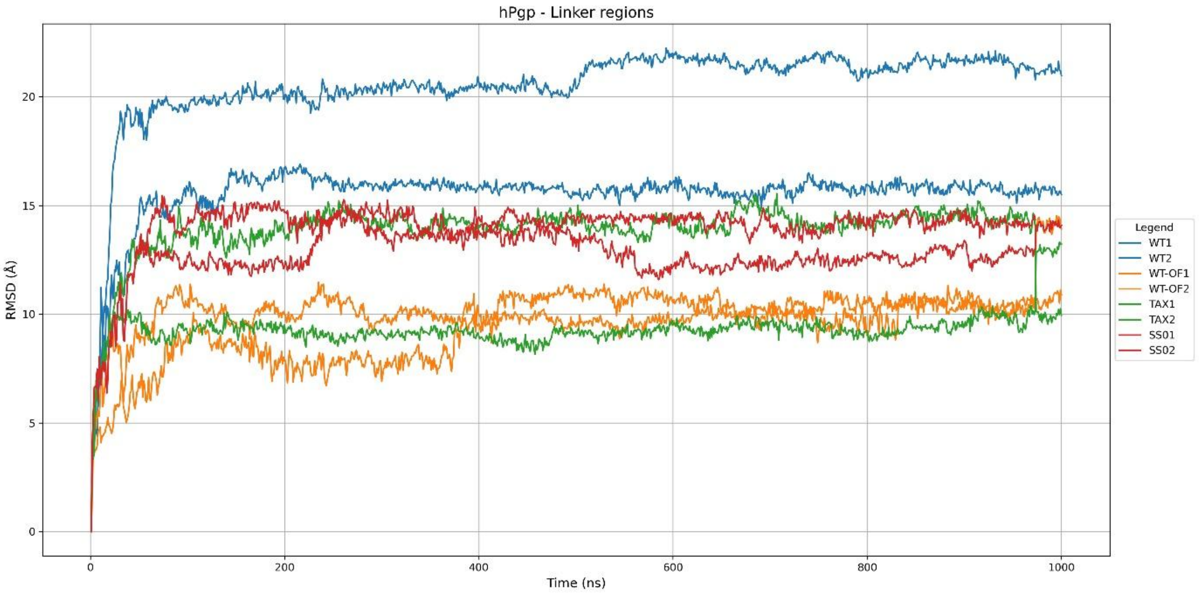


**Figure S12.** Root mean square deviation (RMSD) of the linker regions in selected simulations of the human P-glycoprotein simulations.
